# Water shortage reduces *PHYTOCHROME INTERACTING FACTOR 4, 5* and *3* expression and shade avoidance in Arabidopsis

**DOI:** 10.1101/2022.12.02.518848

**Authors:** Mariana Semmoloni, Cecilia Costigliolo Rojas, Yan Yan, Xiaofeng Cao, Jorge J. Casal

## Abstract

In agricultural crops, forests and grasslands, water deficit often occurs in the presence of cues from neighbouring vegetation. However, most studies have addressed separately the mechanisms of plant growth responses to these two aspects of the environment. Here we show that transferring *Arabidopsis thaliana* seedlings to agar containing polyethylene glycol (PEG) to restrict water availability reduces hypocotyl growth responses to shade without simultaneous affecting cotyledon expansion or its response to shade. Water restriction diminished the activity of the *PHYTOCHROME INTERACTING FACTOR 4 (PIF4), PIF5, PIF3* and *PIF3-LIKE 1* gene promoters, particularly in seedlings exposed to simulated shade. The response of *PIF4* expression to PEG required the presence of its positive morning regulators *CIRCADIAN CLOCK ASSOCIATED 1 (CCA1)* and *LATE ELONGATED HYPOCOTYL (LHY)*, which also reduced their expression in response to PEG. Water restriction diminished the nuclear abundance of PIF4 in hypocotyl cells only in the seedlings exposed to shade. In addition to the changes in *PIF4* levels, post-transcriptional processes also contributed to the response to PEG. Hypocotyl growth showed significant triple interaction among water availability, shade and the presence of PIF4, PIF5 and PIF3. Collectively, these results unveil PIFs as a hub that interlinks shade and drought information to control growth.

## INTRODUCTION

The shade imposed by neighbours within plant canopies entails a reduction in the radiation for photosynthesis available per individual and generates light cues perceived by sensory receptors. These cues include an overall reduction in irradiance from UV-B to far-red, and changes in spectral distribution of the radiation because tissues containing photosynthetic pigments absorb more strongly in red and blue light than in green light and particularly far-red light (Casal, 2013; Ballaré and Pierik, 2017; Roig-Villanova and Martínez-García, 2016). Upon perception by the sensory receptors, these changes in the light microenvironment initiate signalling cascades controlling plant architecture. For instance, light cues from neighbours enhance stem growth and plant stature, a typical shade-avoidance response.

Under sunlight conditions, the activity of the photo-sensory receptors phytochrome B (phyB) and cryptochrome 1 (cry1) repress the growth of hypocotyl in *Arabidopsis thaliana* (Sellaro *et al*., 2010). Light cues from neighbours reduce the activity of these receptors releasing hypocotyl growth from their inhibitory effect. This promotion of hypocotyl growth involves an early burst of auxin synthesis in the cotyledons and its transport to the hypocotyl (Tao *et al*., 2008; Procko *et al*., 2014) and a subsequent increase in auxin sensitivity (Bou-Torrent *et al*., 2014; de Wit *et al*., 2015; Pucciariello *et al*., 2018). PHYTOCHROME INTERACTING FACTOR 4 (PIF4), PIF5, PIF3 and PIF7 are a set of bHLH transcription factors of fundamental importance in the promotion of hypocotyl growth in response to the cues from neighbours (Lorrain *et al*., 2008; Li *et al*., 2012; Sellaro *et al*., 2012; Leivar *et al*., 2012). They activate the expression of a large set of genes in the cotyledons and/or hypocotyl (Kohnen *et al*., 2016), including auxin synthesis *YUCCA* genes (Hornitschek *et al*., 2012; Li *et al*., 2012), auxin transport genes such as *PIN-FORMED 3 (PIN3)* (Keuskamp *et al*., 2010) and auxin signalling genes such as *INDOLE-3-ACETIC ACID INDUCIBLE 19 (IAA19)* (Pucciariello *et al*., 2018). These genes, among many others, mediate the hypocotyl growth promotion. In addition, they induce the expression of regulatory genes that adjust the magnitude of the shade avoidance response such as *LONG HYPOCOTYL IN FAR-RED (HFR1)* and *PHYTOCHROME INTERACTING FACTOR 3-LIKE 1 (PIL1)* (Salter *et al*., 2003; Sessa *et al*., 2005; Hornitschek *et al*., 2009; Hao *et al*., 2012; Roig-Villanova *et al*., 2006; Li *et al*., 2014). PIFs physically interact with phyB, which reduces their activity by diverse molecular mechanisms, including the facilitation of their phosphorylation followed by ubiquitination and proteasome degradation, their decreased transactivation of target genes and/or their retention in the cytosol (Pham *et al*., 2018). Cry1 also interacts with PIF4, to repress its transcription activity (Pedmale *et al*., 2016; Ma *et al*., 2016). Thus, neighbour cues increase the activity of PIFs to mediate the shade-avoidance response (Lorrain *et al*., 2008; Li *et al*., 2012; Leivar *et al*., 2012; Huang *et al*., 2018; Pucciariello *et al*., 2018).

Multiple simultaneous stresses can affect plants in the field (Zandalinas and Mittler, 2022); i.e., plants can simultaneously experience shade and drought. Water deficit lowers growth rates mainly through reduced cell-wall loosening rather than by decreasing turgor pressure (Bogoslavsky and Neumann, 1998; Feng *et al*., 2016; Schopfer, 2006; Claeys and Inzé, 2013). Water deficit reduces cell-wall acidification (Van Volkenburgh and Boyer, 1985; Bogoslavsky and Neumann, 1998), which is required for the cell-wall loosening activity of expansins (Cosgrove, 2005). When transferred to agar containing polyethylene glycol (PEG) to lower the water potential of the substrate, the hypocotyl of *Arabidopsis thaliana* shows reduced growth (Van Der Weele *et al*., 2000). However, *A. thaliana* seedlings rapidly accumulate proline and other solutes via abscisic acid (ABA)-dependent and independent mechanisms to lower cell osmotic potential and restore turgor pressure (Chiang and Dandekar, 1995; Verslues and Bray, 2006; Ozfidan *et al*., 2013). How plant cells perceive a deteriorated water status to control cell-wall extensibility remains unclear but the mechanism could involve the wall-plasma membrane interface (Vaahtera *et al*., 2019; Haswell and Verslues, 2015). Since both shade and water restriction affect cell wall extensibility (Sasidharan *et al*., 2010; Feng *et al*., 2016), the output of combined shade and drought on growth is difficult to predict.

In rice, drought reduces the expression of the PIF-like *OsPIL1* gene (Todaka *et al*., 2012), which promotes stem growth (Todaka *et al*., 2012). In transgenic plants of *A. thaliana*, the expression of *OsPIL1* counteracted the reduced hypocotyl growth caused by the overexpression of DEHYDRATION RESPONSE ELEMENT B1A (DREB1A) used to improve drought resistance (Kudo *et al*., 2017). Maize PIFs enhance plant stature and the response to neighbour cues (Wu *et al*., 2019) but in contrast to *OsPIL1, ZmPIF3* and *ZmPIF1* increase their expression in response to PEG (Gao *et al*., 2015; Gao *et al*., 2018). Furthermore, In contrast to Arabidopsis and maize PIFs (Pham *et al*., 2018; Wu *et al*., 2019), *OsPIL1* does not interact with phytochrome (Todaka *et al*., 2012) and is more stable than PIF4 in the light (Kudo *et al*., 2017). Therefore, it is not clear whether PIFs can represent a point of integration of information from shade and water conditions. Since the closest *A thaliana* orthologs of OsPIL1 are PIF4 and PIF5 (Nakamura *et al*., 2007), we investigated whether water restriction affects the expression of *PIF4, PIF5* and on a second step *PIF3* (the founder family member) and the consequences of these effects on the shade-avoidance response of *A. thaliana* seedlings.

## RESULTS

### Water restriction reduces shade avoidance

We transferred the seedlings grown under control conditions in the laboratory to agar containing the non-permeable high-molecular-weight osmolyte PEG to reduce water availability (Osmolovskaya *et al*., 2018; Dubois and Inzé, 2020). The PEG treatment started one day before the simulated shade treatment to avoid the early impact of the osmotic stress (Van Der Weele *et al*., 2000; Dubois and Inzé, 2020). Then, we measured hypocotyl growth during the first 9 hours of shade treatment, when the initial molecular events of shade avoidance take place. Water restriction reduced the magnitude of the hypocotyl-growth response of *A. thaliana* seedlings to simulated shade (Col-0 wild type in Fig. 1a).

**Figure 1.**
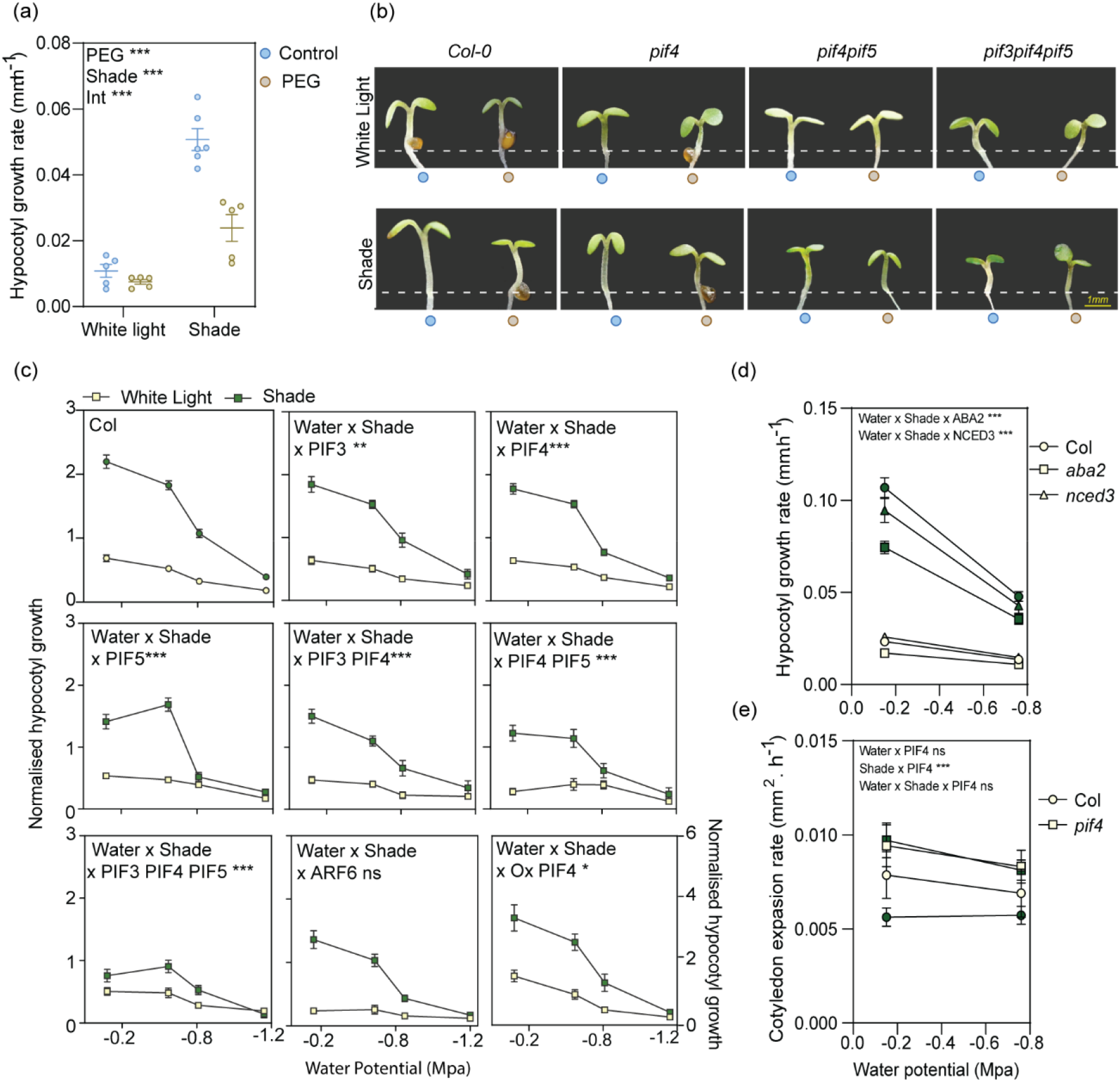
The hypocotyl growth response to water availability under shade requires PIF3, PIF4 and PIF5. (a) Water restriction reduces the shade avoidance response of hypocotyl growth in *A. thaliana*. Data are mean rates of growth (0-9 h of the shade treatment) ±SE and individual values of six replicate boxes (ten seedlings per genotype and per box). The significance of the main effects and interaction is indicated. (b) Photographs of representative seedlings during the second day of shade treatment. Hypocotyl lengths correspond to the average of three replicate boxes with at least three seedlings per genotype and box. (c-d) Hypocotyl growth in the wild type and *pif3, pif4* and *pif5* simple and multiple mutants, the PIF4 overexpressor and the *arf6* mutant (c) or in the *aba2* and *nced3* mutants (d) under white light or shade and different water potentials caused by PEG. Each datum point shows the mean rates of growth (0-9 h of the shade treatment) ±SE of more than 15 seedlings. The significance of the triple interaction among water availability, shade and genotype is indicated. (e) Cotyledon expansion in the wild type and the *pif4* mutant as affected by light or shade and water or PEG. Each datum point shows the mean rates of growth (0-9 h of the shade treatment) ±SE of more than 15 seedlings. The significance of all main effects and possible interactions is indicated (to highlight the lack of effect of PEG). See Experimental Procedures, Statistics, for further details. ***, P <0.001; ns, not significant.

To investigate whether the reduction in the magnitude of shade avoidance by water restriction is a general phenomenon we conducted a series of glasshouse experiments. We generated micro-swards of *A. thaliana* or *Solanum lycopersicon* plants grown in the glasshouse at different population densities, with or without restriction of water availability. To reduce water availability, we used PEG (*A. thaliana*) or restricted irrigation to maintain 30% of the field capacity to reproduce the gradual generation of water deficit that occurs in the field in response to drought (*S. lycopersicon*). Plant density increased the leaf-area index (LAI, leaf area per unit substrate area), enhancing the strength of neighbour cues (Casal, 2013). Hypocotyl growth in *A. thaliana* seedlings and stem growth in *S. lycopersicon* increased with LAI, and PEG reduced the magnitude of this response (Fig. S1a-b). We then used *S. lycopersicon* or *Pisum sativum* grown under sunlight with or without supplementary far-red light to simulate the light reflected by neighbour leaves (Ballaré *et al*., 1987), and reduced water availability by restricting irrigation to maintain 30% of the field capacity. We observed a significant interaction between water availability and supplementary far-red light, indicating that the effects of water restriction were stronger under simulated shade than under sunlight conditions (Fig. S1d, f). Taken together, these results indicate that water restriction reduces the magnitude of shade avoidance responses.

### The response to water availability requires PIF3, PIF4 and PIF5

Taking into account the effects of *OsPIL1* in the response to drought in rice (Todaka *et al*., 2012), we investigated the role of its closest orthologues in *A. thaliana*, i.e., PIF4, PIF5 and expanded the analysis to PIF3, the founder member of the family. Figure 1b illustrates that the magnitude of the response to water availability decreased progressively in the *pif4, pif4 pif5* and *pif3 pif4 pif5* mutants. Figure 1c plots the growth of single and multiple *pif3, pif4* and *pif5* mutants against the water potential of the substrate containing different concentrations of PEG. In all the cases we observed a significant triple interaction among simulated shade, water availability and genotype, indicating that shade has larger effects when water is available and PIF3, PIF4 and PIF5 are present. Consistently with the loss-of-function analysis, overexpression of PIF4 increased the impact of water availability (Fig. 1c). Contrary to the *pif4, pif5* and *pif3* mutants, the *auxin response factor 6 (arf6)* mutation, affecting another transcription factor, reduced hypocotyl growth (Reed *et al*., 2018) but did not diminished the impact of water availability under shade (Fig. 1c).

One of the typical consequences of water deficit is the increase of abscicic acid (ABA) levels (Chen *et al*., 2020; Christmann *et al*., 2005). If ABA had mediated the inhibition of hypocotyl growth caused by PEG, the *aba deficient 2 (aba2)* and/or *nine-cis-epoxycarotenoid dioxygenase 3 (nedc3)* mutations, deficient in ABA synthesis should alleviate the inhibition caused by PEG, increasing hypocotyl growth under PEG and shade. We reject this hypothesis because none of these mutations increased hypocotyl growth (Fig. 1d). On the contrary, both mutations caused a small but statistically significant reduction of hypocotyl growth under shade and high water availability. The latter observation is consistent with the previously recognised role of endogenous ABA in the promotion of vegetative growth (Barrero *et al*., 2005).

### PEG has no rapid effects on hypocotyl growth

To investigate whether water restriction causes a generalised reduction in shoot growth, we measured the rate of expansion of the cotyledons within the same time frame used for hypocotyl growth. The PEG treatment did not reduce cotyledon expansion in any of the conditions (Fig. 1e). Shade did reduce cotyledon expansion in a PIF4-dependent manner (Fig. 1e). Although water restriction had no rapid effects on cotyledon expansion, it was effective later and reduced final cotyledon area measured 3.5 d after the beginning of PEG treatments (mean ±SE of five replicate boxes of seedlings, cm^2^, white light, control= 0.66 ±0.04; PEG= 0.46 ±0.01; shade, control= 0.58 ±0,05, PEG= 0.47 ±0,02; PEG: P= 0.0005). Taken together, these observations demonstrate that there is no generalised growth reduction; rather, a rapid and specific hypocotyl growth response to water restriction before cotyledon expansion becomes affected.

### Water restriction reduces the expression of shade-avoidance gene markers

To investigate whether drought effects on growth responses to shade involve the molecular machinery, we also analysed the activities of the *PIL1, IAA19* and *HFR1* promoter fused to the LUC reporter and of the *PIN3* promoter fused to GUS, as these are four classical molecular markers of shade avoidance and targets of PIFs (Pedmale *et al*., 2016; Kohnen *et al*., 2016; Oh *et al*., 2014). PEG significantly reduced the promotion by shade of the bioluminescence driven by *PIL1* and *IAA19*, without significantly affecting the response driven by *HFR1* (note the significant interaction between PEG and shade in Fig. 2a). In the case of *PIN3*, PEG reduced the expression to similar levels under white light and shade (Fig. 2b). Thus, drought not only affects growth responses to shade but also the expression of selected gene markers.

**Figure 2.**
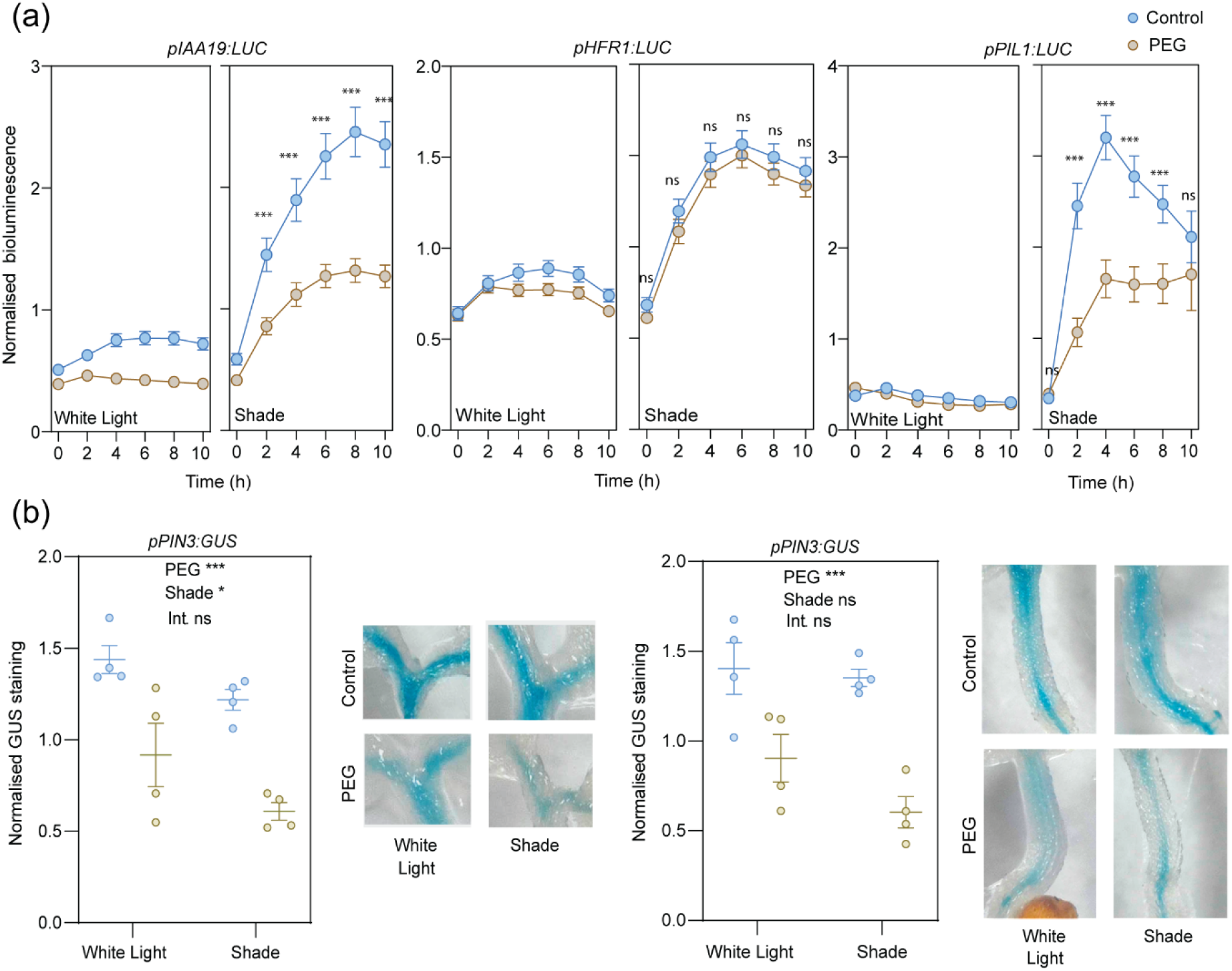
Water restriction reduces the promoter activity of selected shade-avoidance gene markers. (a) Promoter activities analysed by luminometry in seedlings bearing the *pIAA19-LUC, pHFR1-LUC* or *pPIL1-LUC* transgenes. Data points are means ±SE of 70 *(IAA19)*, 72 *(HFR1)* or 28 *(PIL1)* seedlings. The significance of the interaction between PEG and shade is indicated for each time point. (b) Promoter activities in the cotyledons and the hypocotyl analysed by the quantification of GUS staining in seedling bearing the *pPIN3:GUS* transgene. Mean ±SE and individual values of four replicate boxes of seedlings (eight seedlings per box) and representative seedlings. The significance of the main effects and interaction is indicated. *, P <0.05; **, P <0.01; ***, P <0.001; ns, not significant.

### Drought reduces *PIF3, PIF4* and *PIF5* expression

The above results admit two explanations: Either PIFs are simply a condition for the response to water availability or PIFs are part of the response to water availability. If they transmit the water availability signal, PEG should affect some feature of the dynamics of PIFs. Figure 3a-b shows that drought reduced the activity of the *PIF3, PIF4* and *PIF5* promoters fused to the GUS reporter both in the cotyledons and in the hypocotyls. In the cotyledons, PEG affected these gene promoters under simulated shade and under white light conditions (Fig. 3a). In the hypocotyl, *PIF4* responder to PEG under both conditions but *PIF3* and *PIF5* only under shade (Fig. 3b).

**Figure 3.**
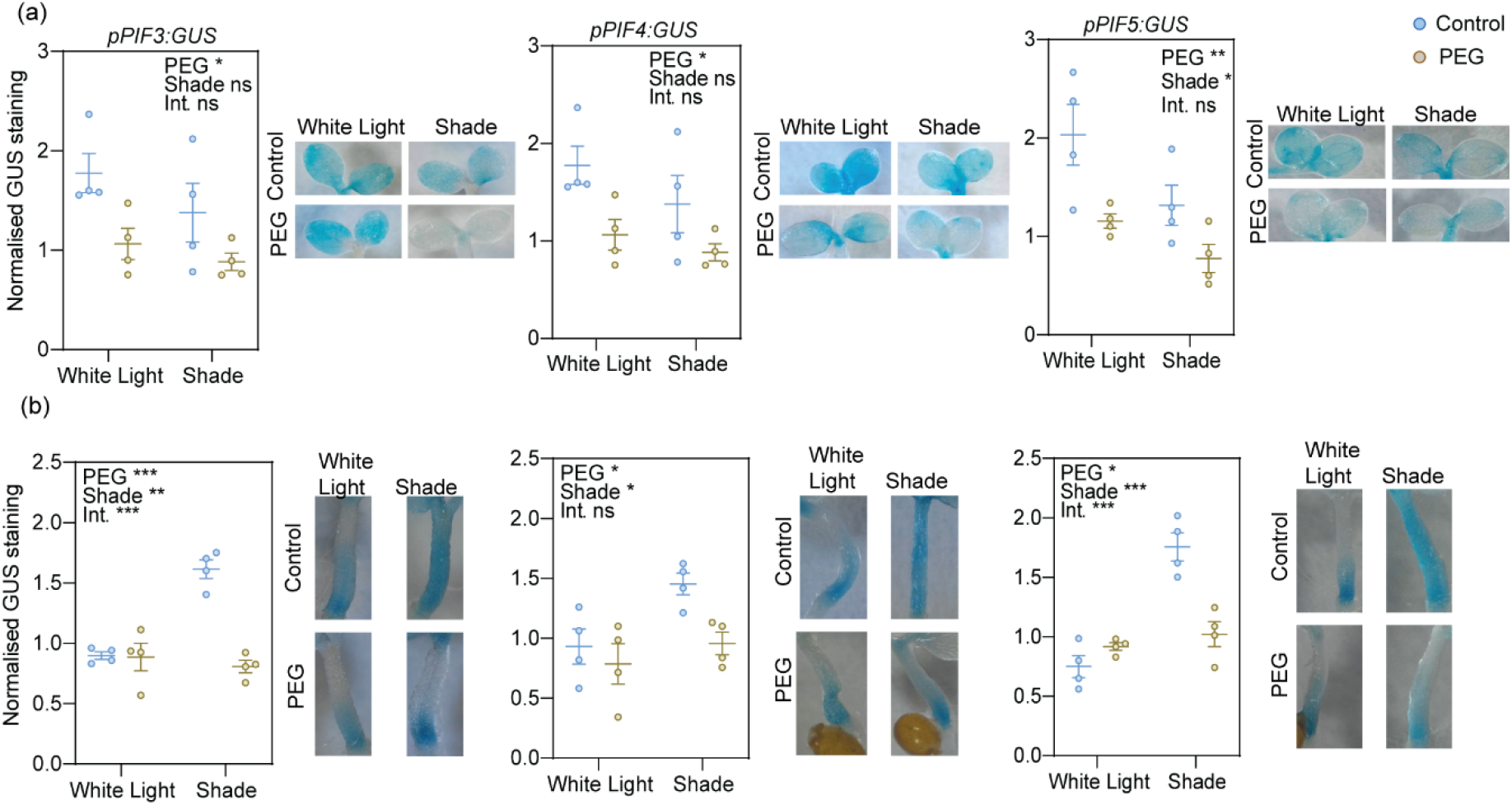
Water restriction reduces *PIF3, PIF4* and *PIF5* promoter activities. Promoter activities in the cotyledons (a) and the hypocotyl (b) analysed by the quantification of GUS staining in seedling bearing the *pPIF3:GUS, pPIF4:GUS* or *pPIF5:GUS* transgene. Mean ±SE and individual values of four replicate boxes of seedlings (six seedlings per box) and representative seedlings. The significance of the main effects and interaction is indicated. *, P <0.05; **, P <0.01; ***, P <0.001; ns, not significant.

Figure 4a, shows the kinetics of *PIF4* promoter activity in luminometer experiments. Since the seedlings are recorded from above, bioluminescence reflects promoter activity mainly in the cotyledons. As observed for the experiments with the GUS reporter, there was a significant effect of PEG but no interaction (the fall in activity caused by PEG was similar under light or shade).

**Figure 4.**
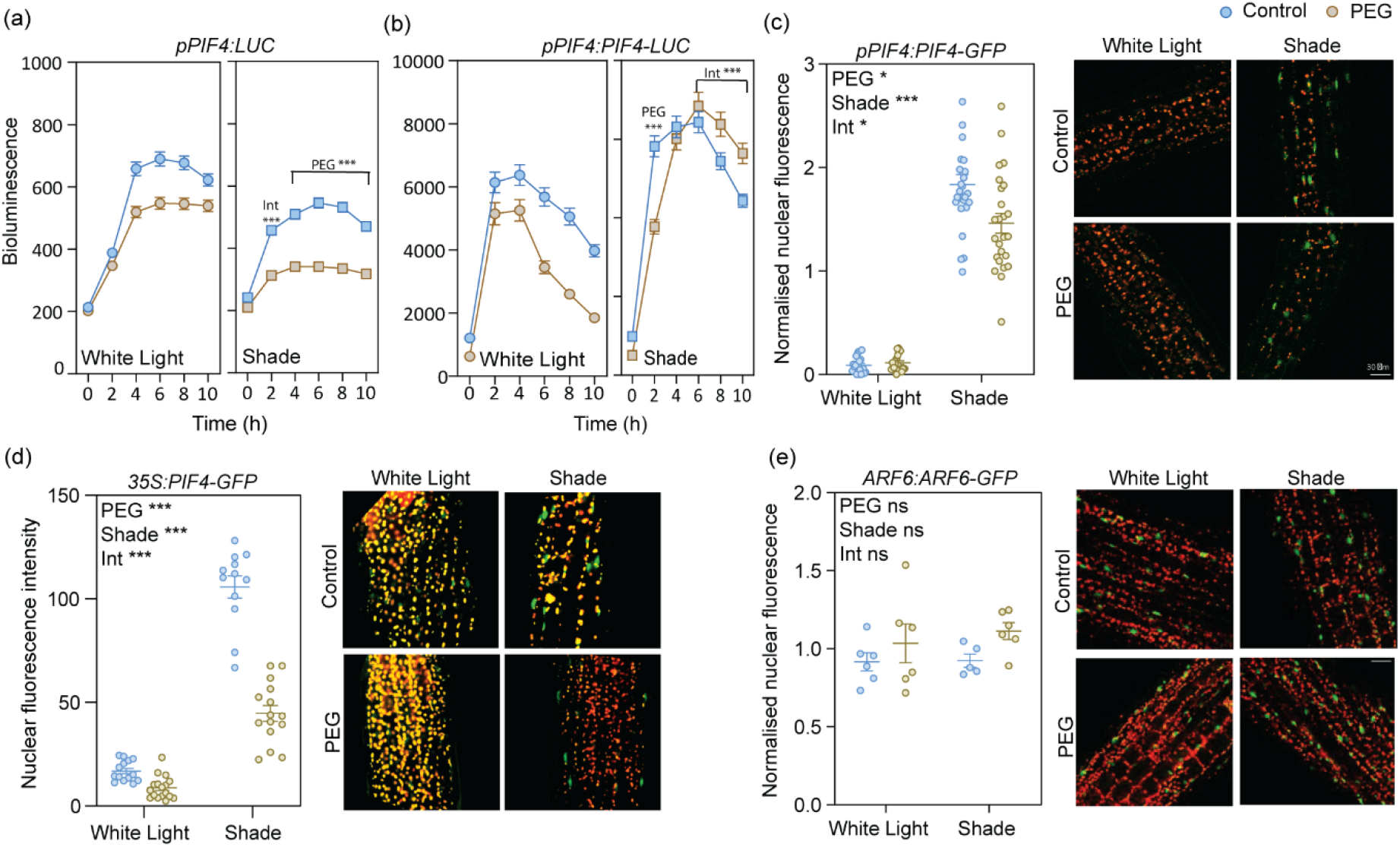
Water restriction reduces PIF4 protein abundance. (a-b) Time course of *PIF4* promoter activity (a) and PIF4 protein abundance (b) analysed by luminometry in seedlings bearing the *pPIF4:LUC* (a) or the *pPIF4:PIF4-LUC* transgene (b). Data points are means ±SE of 120 (a) or 182 seedlings. (c-e) PIF4 (c-d) and ARF6 (e) nuclear abundance in the hypocotyl cells analysed by confocal microscopy in seedlings bearing the *pPIF4:PIF4-GFP* (c) or the *p35S:PIF4-GFP* (d) or *pARF6:ARF6-GFP* (e) transgene. Mean ±SE and individual values of at least 15 (PIF4) or five (AEF6) replicate boxes of seedlings (three seedlings per box) and representative seedlings. *, P <0.05; **, P <0.01; ***, P <0.001; ns, not significant.

### Drought reduces PIF4 abundance

To investigate whether the levels of PIF4 protein reflect the increased expression of *PIF4*, we used lines expressing PIF4 under the control of its own promoter and fused to a bioluminescent reporter. The cotyledon signal dominates the results observed in luminometer experiments. Under white light, PEG reduced the abundance of PIF4 throughout the photoperiod (Fig. 4b). Under shade, PEG reduced the abundance of PIF4 in the morning and increased PIF4 levels during the afternoon (Fig. 4b). The effects of PEG on *PIF4* promoter activity account (at least in principle) for the effects of PEG on PIF4 protein levels under white light and the morning effects of PEG under shade (compare Fig. 4a and b), but the afternoon effects under shade indicate a post-transcriptional control.

We also used lines expressing PIF4 under the control of its own promoter and fused to fluorescent reporter. Confocal microscopy revealed that water restriction reduced the abundance of PIF4 in the hypocotyl (Fig. 4c). The PEG treatment also reduced nuclear fluorescence driven by the *PIF4* transgene under the control of a constitutive promoter (Fig. 4d), indicating the occurrence of post-transcriptional effects in addition to the effects on *PIF4* promoter activity shown above (Fig. 3, 4a).

### Water restriction reduces the expression of *LHY* and *CCA1*

Since CIRCADIAN CLOCK ASSOCIATED 1 (CCA1) and LATE ELONGATED HYPOCOTYL (LHY) bind to the *PIF4* promoter to enhance its activity during the morning (Sun *et al*., 2019), PEG reduces the expression of *CCA1* (Litthauer *et al*., 2018) and suboptimal watering reduces the expression of *LHY* (Dubois *et al*., 2017), we investigated whether CCA1 and LHY mediate the effect of water restriction on the activity of the *PIF4* promoter. Luminometer readings showed and reduced *LHY* and *CCA1* promoter activity in response to PEG (Fig. 5a). As a reference we included *TIMING OF CAB EXPRESSION 1 (TOC1)*, because ABA produces a transient increase in *TOC1* promoter activity followed by a decrease (Legnaioli *et al*., 2009) and restricted watering reduces the expression of *TOC1* (Dubois *et al*., 2017). Consistently with the latter report, PEG reduced the activity of the *TOC1* promoter in the afternoon (Fig. 5a). However, arguing against a direct role of TOC1, PEG had no significant effects on *TOC1* under shade during the morning, a condition where it strongly affects *PIF4* promoter activity (Fig. 5a). The double *lhy cca1* mutant showed reduced *PIF4* expression and no significant response to PEG (Fig. 5b). Taken together, these observations are consistent with a role of LHY and CCA1 in the control of *PIF4* expression by PEG. Contrary to PEG, the addition of exogenous ABA enhanced *PIF4* promoter activity (Fig. 5c), which is consistent with previous observations (Liang *et al*., 2020).

**Fig. 5.**
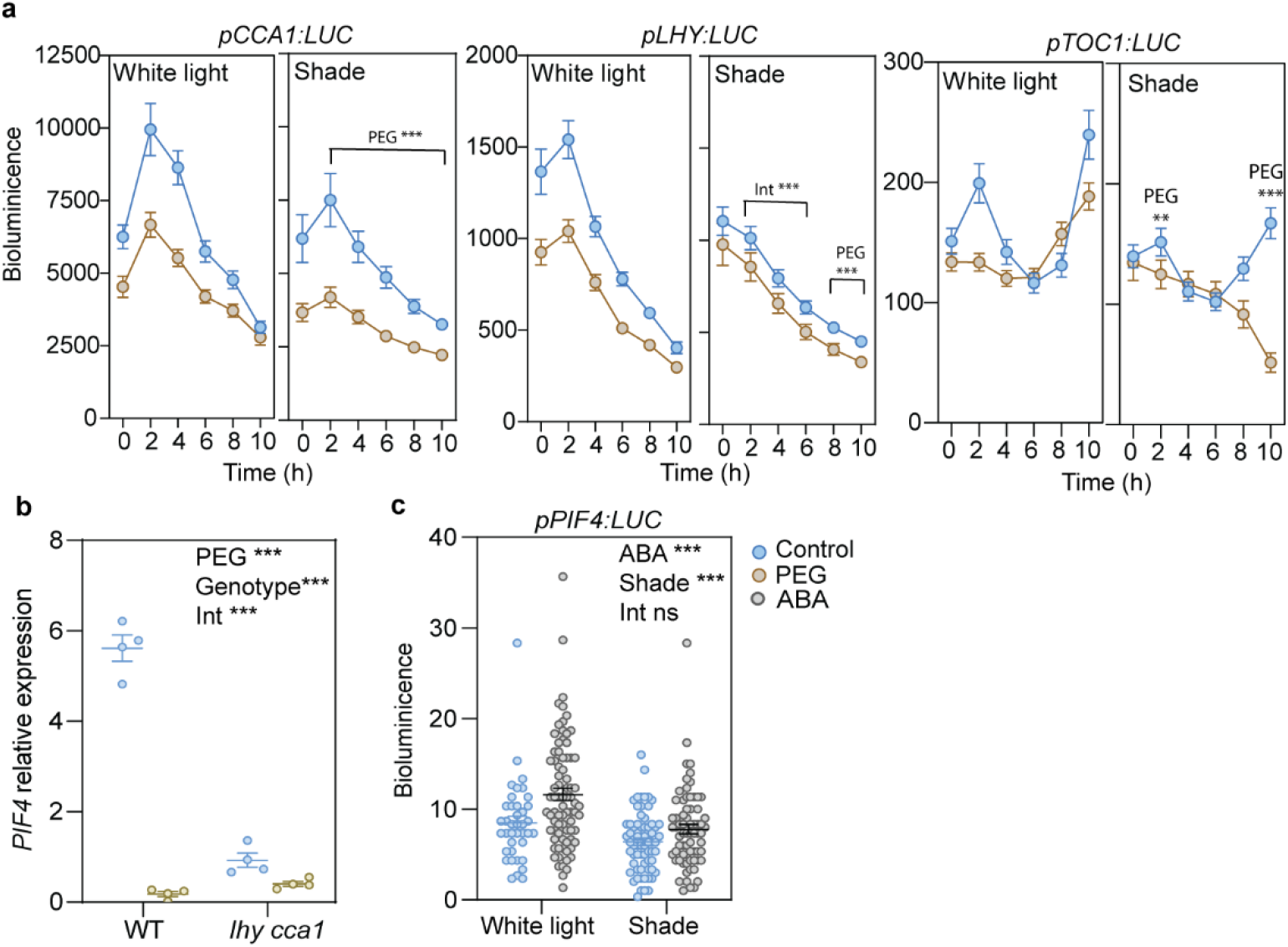
Drought reduces *CCA1* and *LHY* promoter activities to control *PIF4* expression. (a) PEG reduces *CCA1, LHY1* and *TOC1* promoter activity analysed by luminometry in seedlings bearing the *pCCA1:LUC, pLHY1:LUC* or *pTOC1:LUC* transgene expression. Data points are means ±SE of at least 24 seedlings. (b) PEG reduces the expression of *PIF4* analysed by real-time PCR, in a CCA1-LHY-dependent manner. Mean ±SE and individual values of four replicate boxes of seedlings. (c) Exogenous ABA does not reduce *PIF4* promoter activity analysed by luminometry in seedlings bearing the *pPIF4:LUC* transgene. Data points are means ±SE of at least 30 seedlings.

## DISCUSSION

We show that low water availability impairs shade avoidance in *A. thaliana, S. lycopersicum* and *P. sativum* (Fig. 1a, S1). This effect is specific, not the consequence of an overall contraction of the growth rates, because water restriction did not affect cotyledon expansion within the same time frame when it severely reduced hypocotyl growth in *A. thaliana* (Fig. 1e).

Addition of PEG to the growth substrate diminished the activity of the *PIF4, PIF5* and *PIF3* promoters particularly under shade (Fig. 3), which is consistent with the reduced *PIF4* expression observed in *A. thaliana* rosettes exposed to drought (Kudo *et al*., 2019). PEG also lowered the abundance of the PIF4 protein under shade (Fig. 4) and the activity of *IAA19, PIN3* and *PIL1* promoters (Fig. 2), which are targets of PIFs (Pedmale *et al*., 2016; Kohnen *et al*., 2016; Oh *et al*., 2014). Upstream of PIF4, PEG reduced the expression of *CCA1* and *LHY1* (see also Litthauer *et al*., 2018; Dubois *et al*., 2017), which are required both to promote *PIF4* expression during the morning (Sun *et al*., 2019) and for the response of *PIF4* expression to PEG (Fig. 5a-b). Overexpression of PIF4 did not eliminate the interaction between PEG, shade and PIF4 on hypocotyl growth (Fig. 1c), suggesting that water restriction has effects beyond enhancing the expression of *PIF4*. In fact, PEG reduced the nuclear abundance of PIF4 driven by a constitutive promoter (Fig. 4d), indicating the occurrence of post-transcriptional effects on PIF4. In turn, the *pif4*, *pif5* and/or *pif3* mutations reduced the magnitude of the effect of PEG on hypocotyl growth particularly in the seedlings exposed to shade (Fig. 1b-c). Taken together, these results indicate that water restriction diminishes the activity of PIF4, PIF5 and PIF3, which in the case of PIF4 involves transcriptional effects mediated by CCA1 and LHY1 as well as post-transcriptional effects, and consequently restrains shade avoidance. This model does not exclude putative effects of water restriction on the activity of other components of the shade avoidance network.

PIFs have a dual action in the promotion of hypocotyl growth in response to neighbour cues. One function is to promote a rapid increase in auxin synthesis in the cotyledons (Hornitschek *et al*., 2012; Li *et al*., 2012), which is transported to the hypocotyl (Tao *et al*., 2008; Procko *et al*., 2014). The other action is to promote growth locally in the hypocotyl itself by enhancing the sensitivity to auxin (Kohnen *et al*., 2016; Pucciariello *et al*., 2018). In the cotyledon, which is the site of perception of neighbour cues (Procko *et al*., 2014), *PIF4* and *PIF5* responded to PEG even in seedlings exposed to white light (Fig. 3a). Luminometer readings, dominated by the signal produced by the cotyledons, also showed reduced PIF4 protein levels in response to PEG under white light (Fig. 4b). Therefore, in plants suffering water restriction, this organ appears preconditioned to favour a weaker response to shade. In the hypocotyl, PEG reduced the expression of *PIF5* and *PIF3* and the abundance of PIF4 only in the seedlings exposed to shade (Fig. 3b, 4c). This pattern is important to account for the impaired shade avoidance response caused by PEG, i.e., not just a general reduction of hypocotyl growth but a stronger curtailment of growth under shade. The observation also supports the concept that some molecular responses occur only when different stresses are combined (Zandalinas and Mittler, 2022), as it is the case here for water restriction and shade.

PIF4 promotes hypocotyl growth but inhibits cotyledon expansion (Huq and Quail, 2002). The PIF4-mediated inhibition of cotyledon expansion is particularly important under shade (Costigliolo Rojas *et al*., 2022). Interestingly, under shade, cotyledon PIF4 protein levels observed in luminometer experiments decreased in the morning and increased in the afternoon in response to PEG compared to the control (Fig. 4b). The enhanced afternoon PIF4 abundance in response to PEG could help to neutralise the effect of the morning drop in PIF4 (required to reduce the hypocotyl growth response to shade) and maintain the PIF4-mediated shade inhibition of cotyledon expansion in the presence of water restriction (Fig. 1e).

Salinity also reduces shade avoidance but, contrary to the PEG effects reported here, salinity does not affect PIF4 or PIF5 abundance or *PIF4* or *PIF5* expression (Hayes *et al*., 2019). Furthermore, whereas the salinity effect on shade avoidance depended on ABA signalling (Hayes *et al*., 2019), the response to PEG did not (Fig. 1d, 5c). It is important to emphasise that, in addition to water restriction, salinity involves competition between ions and toxicity effects. Toxic effects on extension growth are rapid as they can be observed in less than 2 h after the application of NaCl (Veselov *et al*., 2009). Therefore, toxic effects of salinity might be responsible for the reported effects of NaCl on shade avoidance (Hayes *et al*., 2019), accounting for the mechanistic differences observed between both responses. Worthy of note, *OsPIL1* also reduced its expression in response to water restriction but not in response to ABA or salt (Todaka *et al*., 2012).

There is an apparent trade-off between shade avoidance and tolerance to drought. At the molecular level, overexpression of *PIF4* (which enhances shade avoidance, Fig 1c), reduces the levels of *DREB1A* overexpression and drought tolerance in double transgenics, when compared to the parent DREB1A overexpressors (Kudo *et al*., 2019). At the canopy level, in crowded environments, plants mutually compete for light but the reduced radiation load under shade and the shelter from wind impact provided by neighbours decrease the rate of transpiration. Thus, nearby vegetation limits light for photosynthesis but it ameliorates water stress by preventing excessive water loss. Therefore, as water status deteriorates, plant-plant interactions may gradually shift from competitive to facilitative (Holmgren *et al*., 1997). Competitive shade-avoidance responses help to overtop neighbours. When water availability is adequate, these responses increase plant fitness (Dudley and Schmitt, 1996) due to enhanced light interception. However, under restricted water availability, shade avoidance is not beneficial because photosynthesis can be limited despite light availability (Marañón *et al*., 2006). Furthermore, overtopping neighbours when water is scarce can be detrimental due to foliage exposure to sunlight (warming) and wind (McGregor *et al*., 2021), and because the shade-avoidance phenotype tends to be more costly in dry microsites in the field (Huber *et al*., 2004). The PIF-mediated control would adjust the growth pattern when plant-plant interactions shift from competitive to facilitative under restricted water availability.

## EXPERIMENTAL PROCEDURES

### Plant Material

Seeds of *A. thaliana* were sown on filter paper placed on top of 5 ml of 0.8% agar 0.5 MS (Murashige and Skoog, 1962) and incubated in darkness at 5°C for 4 d. We used the Columbia-0 line as wild type and the mutants and transgenic lines listed in Table S1.

### Light conditions

Stratified seeds were transferred to white-light photoperiods (10 h) provided by a mixture of fluorescent and halogen lamps providing 90-100 μmol m^-2^ s^-1^ (400-700 nm) and a red / far-red ratio= 1.1 at 20°C. White light control seedlings remained under this condition whereas those exposed to simulated shade were covered with two green acetate filters (no. 089; LEE Filters, Hampshire, UK) to reduce the radiation to 10 μmol m^-2^ s^-1^ (400-700 nm) and the red / far-red ratio to 0.1 (Pacín *et al*., 2013), one hour after the beginning of the photoperiod of day 4.

### PEG treatments

We used PEG to generate water restriction (Osmolovskaya *et al*., 2018; Verslues *et al*., 2006). We added 1.5 ml solutions containing 0 g/l (water control), 250 g/l, 400 g/l or 700 g/l of PEG of a molecular weight of 8000 (BIOFROXX, 25322-68-3) per 1 ml of agar substrate, incubated for 24 h and discarded the excess of solution. In the experiments where we used a single PEG treatment it corresponds to the intermediate concentration (400 g/l). The resulting water potentials (CS52 psychrometric sample chamber, Wescor Inc., Logan, Utah, coupled to a HR-33T Dew Point Microvoltimeter, Wescor Inc., Logan, Utah) were −0.20 (control), −0.56, −0.75 and −1.20 MPa, respectively, and the pH of the agar was 6.0 (pH indicator strips, Merck, Buenos Aires, Argentina). At the beginning of the photoperiod of day 3 (25 h before the beginning of shade treatments), we transferred the filter paper with seedlings from their original agar to agar equilibrated with the above solutions.

### Growth measurements

In hypocotyl growth experiments, after stratification the boxes were turned to place the agar vertically and favour the subsequent imaging of the hypocotyls in seedlings grown against the agar. In cotyledon expansion experiments, the agar remained horizontal throughout the experiment. At the beginning of the photoperiod of day 3, we transferred the filter paper with seedlings from their original agar to agar equilibrated with the PEG (or distilled water) solutions. To measure the hypocotyl elongation rate or cotyledon expansion rate, the photographs of each box were taken (perpendicularly to the agar) with a digital camera (PowerShot; Canon, http://www.canon.com) at the beginning of the shade treatment and at the end of the photoperiod of the fourth day (times 0 and 9 h, respectively). We used Image analysis software (Adobe Photoshop CC) to superimpose the photos corresponding to times 0 and 9 h and measured the increase in hypocotyl length or cotyledon area over that time. The increase was divided by 9 h to calculate the growth rate. To select images from representative seedlings we photographed each box at midday of the second day under shade and measured hypocotyl length with ImageJ.

### GUS staining

Four hours after the beginning of the photoperiod of day 4, we harvested the seedlings in 250 μl of cold 90% acetone (incubation for 20 min), rinsed the material twice with 250 μl miliQ water and incubated it the dark at 37ºC in 1mM X-Gluc staining solution (5-bromo-4-chloro-3-indolyl-β-D-glucuronide, 10% Triton X100, 100 mM sodium phosphate, 5 mM ferricyanide, 5 mM ferrocyanide and 1 M EDTA). The incubation time depended on each reporter line. We successively washed the stained seedlings in alcohol from 5% to 90% in the dark at room temperature to obtain the required transparency of the plant material and took pictures of the seedlings with an Andonstar ADSM302 digital magnifying glass with 3MP image sensor Incorporated. We converted the images to HSB mode and quantified GUS staining in areas of interest using the channel “Saturation” of the ImageJ software (Béziat *et al*., 2017).

### Bioluminescence assays

At the beginning of the photoperiod of day 3, we transferred each seeding to one of the 96 wells of the luminometry plates and 5 h later we added 20 μl of 0.5 mM D-Luciferin to each well. We detected luciferase activity (three successive seconds per well) every 2 h from the beginning to the end of the photoperiod of day 4 with a luminometer Center LB 960 (Berthold).

### Confocal microscopy

We obtained confocal fluorescence images with an LSM710 and LSM510 (Zeiss) laser scanning microscope with a Plan-Apochromat 40×/1.2 objective lens. For chloroplast visualisation, probes were excited with a He-Ne laser (λ = 543 nm) and fluorescence was detected using an LP560 filter. For visualization of GFP/YFP fusion proteins, probes were excited with an Argon laser (λ = 488 nm) and fluorescence was detected using a BP 505-530 filter. The pinhole was closed to obtain a 5 μm optical sectioning. The images were taken from the epidermis (although this may include the first sub-epidermal layer). Confocal fluorescence images were taken 4 h after the application of the shade treatment. We measured the fluorescence of all the nuclei in each image (we show representative images) and divided the sum of these fluorescence values by the number of cells in the image. Therefore, the nuclei with no detectable fluorescence did not contribute to the numerator of the equation but their cells were included in the denominator. To calculate the number of cells we measured the area of five cells and divided the area of the image by the average area of a cell. We performed image analysis in batch with an image segmentation program developed in Icy (http://icy.bioimageanalysis.org/; Sellaro *et al*., 2019)

### Quantitative reverse transcriptase-PCR

We harvested the seedlings in liquid Nitrogen 4 h after the beginning of day 4 (i.e., after 4 h of shade treatments). Total RNA was extracted with Spectrum Plant Total RNA Kit (Sigma-Aldrich) and subjected to a DNAse treatment with RQ1 RNase-Free DNase (Promega). cDNA derived from this RNA was synthesised using MMLV (Thermofisher) and an oligo-dT primer. The synthesised cDNAs were amplified with FastStart Universal SYBRGreen Master (Roche) using the 7500 Real Time PCR System (Applied Biosystems, available from Invitrogen) cycler. The *UBIQUITIN-CONJUGATING ENZYME 2 (UBC2)* gene was used as the normalisation control (Staneloni *et al*., 2009). The primers are listed in Table S2.

### Statistics

To analyse the effects of water or PEG treatments in combination with light or shade in growth GUS staining or confocal experiments (Figs 1a, 2b, 3, 4c-e, 5b), we averaged the data corresponding to all the seedlings of each growth box, which were assigned at random to the different combinations, and used these average values as replicates in factorial ANOVA (Graphpad software). In luminometer experiments (Fig. 2a, 4a-b, 5a, c) we used the seedlings as replicates in factorial ANOVA because each seedling was assigned at random to one well of the luminometer plate. In real-time PCR we used different boxes as biological replicates in factorial ANOVA after averaging the values of the technical replicates.

To calculate the significance of the triple interaction among water availability as affected by PEG, light or shade and wild type or mutant genotypes (Figs 1c-e), we used the growth values of each seedling (because all genotypes were present in each growth box). We used multiple regression analysis (Statistix software) and in some cases we normalised growth rates to the average of the experiment to obtain a more accurate comparison based on data pooled from different experiments as response variable. Water availability, light condition, genotype and all the possible interactions among them were the explanatory variables. As a proxy for water availability, we calculated 1.2 MPa plus the corresponding water potential in MPa (negative values). Thus, for the highest concentration of PEG (water potential= −1.2 MPa) the variable assumed the value cero. The variable light condition assumed de value cero for white light and one for shade. The variable genotype assumed the value cero for the mutant allele and one for the wild-type allele. We used the step-wise model of multiple regression analysis, which from the original set of variables retains only those that significantly contribute to predict the response variable, pruning those variables that provide redundant information.

## ACKNOWLEDGEMENTS

We thank Dr Romina Sellaro (University of Buenos Aires) for instructing M.S. in the use of the luminometer. This work was supported by grants from the University of Buenos Aires (grant no. 20020170100505BA to J.J.C.) and *Agencia Nacional de Promoción Científica y Tecnológica* (grant numbers PICT-2018-01695 to J.J.C and PICT-2019-2019-2252 to C.C.-R.).

## AUTHOR CONTRIBUTIONS

M.S. and C.C.R conducted the experiments, analysed data, provided input to the research and prepared the figures. Y.Y and X.C. provided input to the research. J.J.C conceived and designed the research, analysed data and wrote the manuscript.

## DATA AVAILABILITY STATEMENT

All data are available upon request to the corresponding author.

## SUPPORTING INFORMATION

Additional Supporting Information may be found in the online version of this article.

**Supplementary Table S1.** Mutant and transgenic lines used in this study.

**Supplementary Table S2.** Primers used for real-time PCR.

**Supplementary Figure S1.** Water restriction by polyethylene glycol (PEG) or drought reduces shade avoidance induced by real neighbours or supplementary far-red light in different species.

---

**Figure S1.**
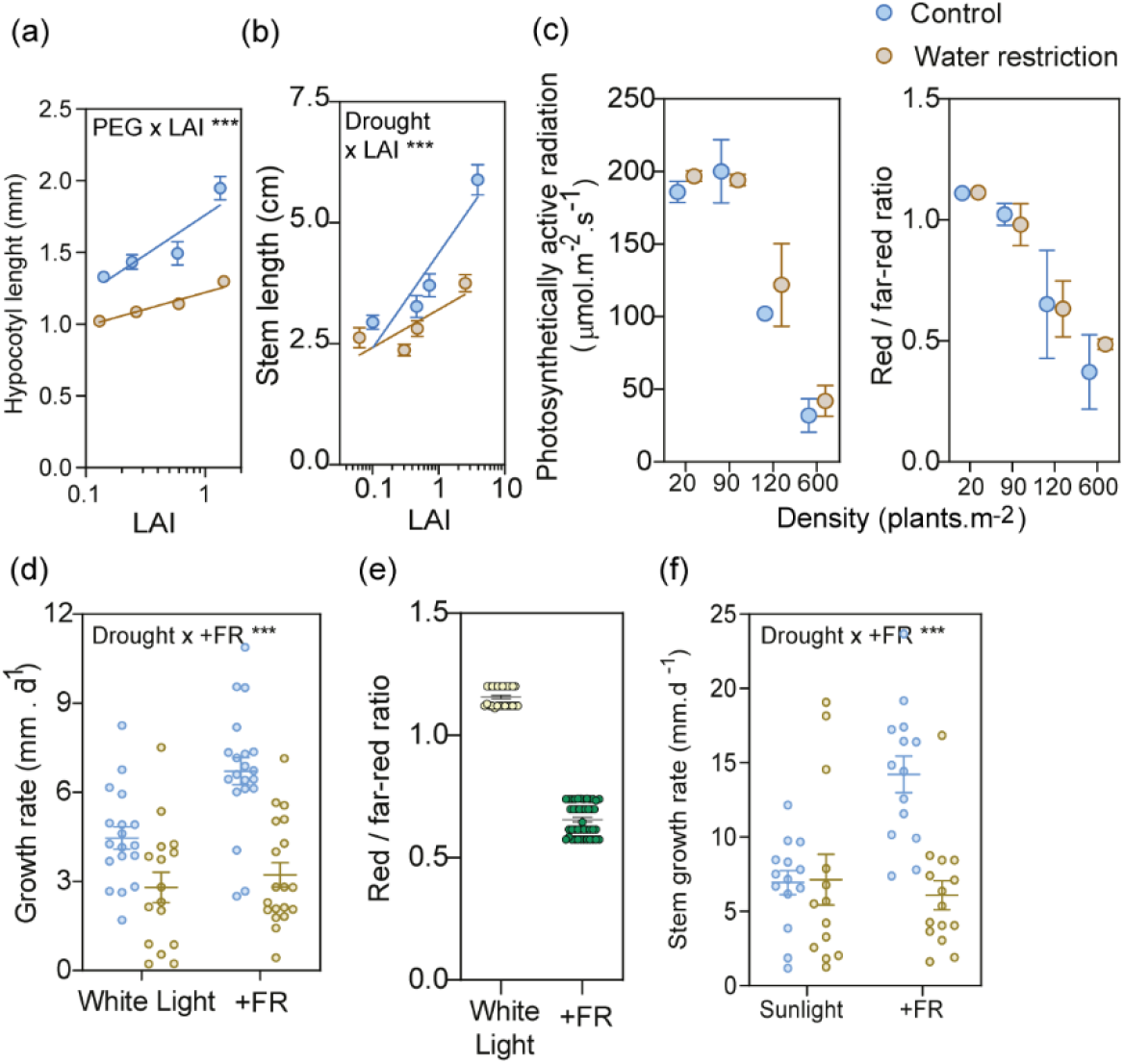
Water restriction by polyethylene glycol (PEG) or drought reduces shade avoidance induced by real neighbours or supplementary far-red light in different species. (a) PEG reduces the promotion of hypocotyl growth caused by neighbours in micro-swards of A. *thaliana*. Different leaf area indexes (LAI) were obtained by varying sowing densities. Means ±SE of at least three replicate boxes of seedlings (at least 12 seedlings measured per box). The interaction between LAI and PEG is significant at P <0.0001. We sow 9, 16, 36, 81 or 145 seeds of wild type *Arabidopsis thaliana* on 2.25 cm^2^ of filter paper placed on top of 0.8% agar-water containing 0.5 Murashige-Skoog (Duchefa). We chilled the seeds and grew the seedlings under controlled conditions for 3 d as described in Experimental Procedures. At the beginning of day 4, we transferred the seedlings to a glasshouse where they remained exposed to natural radiation for another four days. At the end of this period, we measured the area of the cotyledons by using photographs of the seedlings taken from above (still within the box) and image processing software. We also measured the length of the hypocotyl by using photographs taken from the side (after taking the seedlings out of the box) and image processing software. We calculated LAI as the product of area per cotyledon multiplied by two and by the number of seedlings, divided by the sowing area. (b-c) Drought reduces the stem-growth responses of tomato *(Solanum lycopersicon)* to neighbouring vegetation. We arranged pots with one tomato plant at a density of 20, 90, 180 or 600 plants m-2 to achieve the LAI. Length of the stem measured with a calliper after 7 d of treatment (b). Data are means and SE from 20-27 plants (pooled from three experiments) and the interaction between LAI and water restriction is significant at P <0.001. Photosynthetically active radiation (400-700 nm) and red / far-red ratio measured with the remote probe of a SKR 1850 sensor (Skye Instruments) inside the tomato canopies at midday (c). We grew plants of tomato in 330 cm3 plastic pots containing perlite, vermiculite and peat (2-1-1) and watered with Hakaphos Red fertiliser (Compo Expert), in a heated glasshouse under natural radiation. When the plants had their first pair of leaves fully expanded, we suspended the use of fertiliser and randomly assigned half to the control condition (watered to 70-100% of field capacity) and half to the water deficit treatment (watered to 30% of field capacity). The volume of water corresponding to 100% field capacity was the difference in the weight between pots irrigated to saturation (allowed to drain) and the weight of pots containing dry substrate. Plant density treatments started after the water restriction treatment had reached the target percentage of field capacity. (d-e). Drought reduces stem-growth responses of tomato to supplementary far-red light added to natural radiation (to lower the red / far-red ratio). We supplemented natural radiation during the photoperiod (approximately 10 h) with far-red light provided by incandescent lamps (70 W) in combination with a 10-cm thick water filter, a red filter (Lee filter number 26) and three blue filters (Paolini 2031, 2.5 mm thick each one). We measured the length of the stem at the beginning of the far-red light treatment and a day later to obtain the length increment (d). Data are means, SE and individual values of at least 16 plant replicates (pooled from two experiments) and the interaction between the far-red light treatment and water restriction is significant at p < 0.01. Red / far-red ratio measured with the remote probe of a SKR 1850 sensor at the position of the plants, facing the far-red light source, at midday (e). Plant growth and drought treatments as in (b-c). Supplementary far-red light treatments started after the water restriction treatment had reached the target percentage of field capacity. (f) Drought reduces stem-growth responses of pea *(Pisum sativum)* to supplementary far-red light added to natural radiation (to lower the red / far-red ratio). Plant growth and drought treatments as in (b-c). Supplementary far-red light treatment as in (d-e). We measured the length of the stem at the beginning of the far-red light treatment and 2 d later to obtain the length increment. Data are means, SE and individual values of at least 13 plant replicates (pooled from two experiments) and the interaction between the far-red light treatment and water restriction is significant at P < 0.001. The significance of the interaction (Int.) between the effects of water restriction (PER or drought) and neighbour cues (true neighbours or supplementary far-red light) is indicated to support that the effect of water restriction is stronger under actual or simulated shade than under sunlight conditions. **, P <0.01; ***, P <0.001.

**Table S1.**
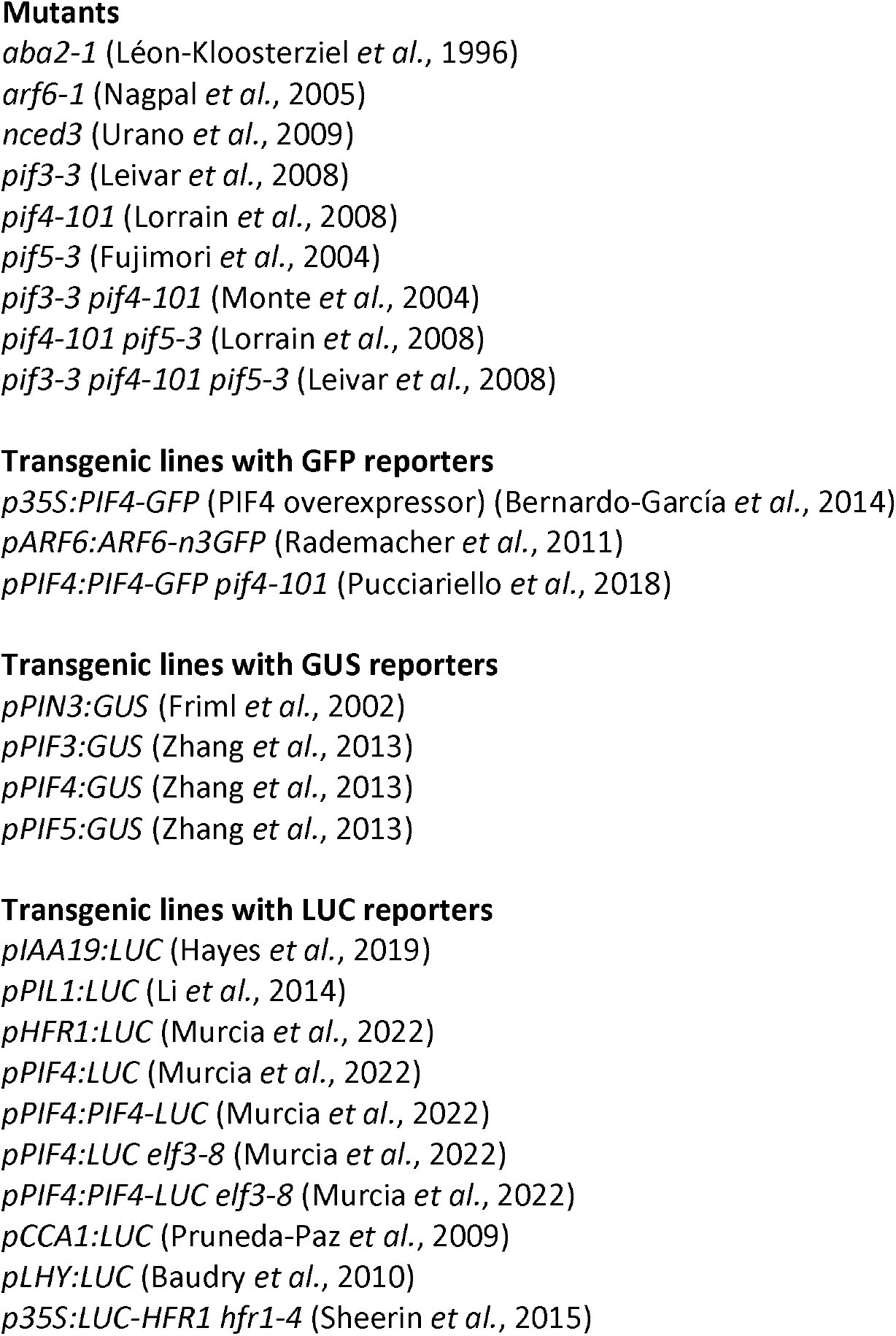
Mutant and transgenic lines used in this study.

**Table S2.**
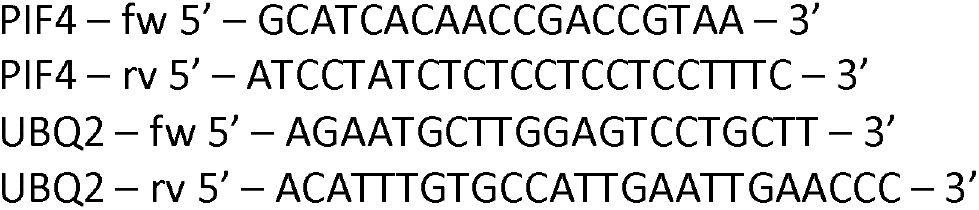
Primers used for real-time PCR.

